# Mathematical modeling of the *Candida albicans* yeast to hyphal transition reveals novel control strategies

**DOI:** 10.1101/2021.01.20.427417

**Authors:** David J. Wooten, Jorge Gómez Tejeda Zañudo, David Murrugarra, Austin M. Perry, Anna Dongari-Bagtzoglou, Reinhard Laubenbacher, Clarissa J. Nobile, Réka Albert

## Abstract

*Candida albicans*, an opportunistic fungal pathogen, is a significant cause of human infections, particularly in immunocompromised individuals. Phenotypic plasticity between two morphological phenotypes, yeast and hyphae, is a key mechanism by which *C. albicans* can thrive in many microenvironments and cause disease in the host. Understanding the decision points and key driver genes controlling this important transition, and how these genes respond to different environmental signals is critical to understanding how *C. albicans* causes infections in the host. Here we build and analyze a Boolean dynamical model of the *C. albicans* yeast to hyphal transition, integrating multiple environmental factors and regulatory mechanisms. We validate the model by a systematic comparison to prior experiments, which led to agreement in 18 out of 22 cases. The discrepancies motivate alternative hypotheses that are testable by follow-up experiments. Analysis of this model revealed two time-constrained windows of opportunity that must be met for the complete transition from the yeast to hyphal phenotype, as well as control strategies that can robustly prevent this transition. We experimentally validate two of these control predictions in *C. albicans* strains lacking the transcription factor *UME6* and the histone deacetylase *HDA1*, respectively. This model will serve as a strong base from which to develop a systems biology understanding of *C. albicans* morphogenesis.

## Introduction

*Candida albicans* is a pleiomorphic, opportunistic fungal pathogen and an important cause of both superficial and systemic infections in humans, particularly in immunocompromised individuals. It is also responsible for 85-95% of all vulvovaginal infections resulting in doctor visits in otherwise healthy patients [1]. *C. albicans* forms biofilms on mucosal surfaces (*e*.*g*., oral, gastrointestinal tract, genitourinary tract, and vaginal) of the host as well as on surfaces of implanted medical devices (e.g., catheters, heart valves, and prosthetics), which are major reservoirs for infections [2,3].

Transitions between the yeast and hyphal phenotypes enable *C. albicans* to adapt to and persist in a wide range of environments. The yeast-form consists of single round cells that grow by forming daughter cells that bud and separate from mother cells. The hyphal form consists of long, multicellular branching tubular structures with parallel-sided walls, where the tips proliferate to elongate the hyphae [4]. *C. albicans* can also form an intermediate filamentous morphology called the pseudohyphal form, which consists of chains of cells with constrictions between mother-daughter cell pairs [4]. Transitioning from the yeast to hyphal phenotype is required for mucosal invasion [2,5] and biofilm formation, which are important mediators of infection [2,3,5,6]. The yeast to hyphal transition is regulated by many well-studied intracellular pathways that respond to external signals such as neutral or alkaline pH (pH > 6), farnesol levels, and temperature. These pathways converge on a handful of key transcription factors, defined as sequence-specific DNA-binding proteins, which regulate the transcription of hyphal-associated genes (HAGs). Epigenetic effects such as histone acetylation events at the promoters of HAGs also play important roles in the regulation of the expression of HAGs. The key negative regulator of the transcription of HAGs is Nrg1. The pattern of expression of Nrg1 and its ability to bind to the promoter region of HAGs determines two phases of the yeast to hyphal transition. Hyphal initiation (the first cell division in the process that forms hyphae) requires a transient downregulation of the Nrg1 protein, whereas hyphal maintenance requires preventing Nrg1 from binding to the promoters of HAGs [7]. External signals initiate the downregulation of *NRG1* transcription, while Nrg1 protein is prevented from binding to the promoters of HAGs by histone deacetylases (HDACs) such as Hda1.

Here we build a Boolean model integrating multiple extracellular signals governing the intracellular regulation of the yeast to hyphal morphological transition. We then use this model to conduct a thorough analysis of phenotype control, considering multiple possible control objectives and side effects, to rank the best and most robust control strategies.

## Results

### Construction of the model

As a starting point to building a model of the intracellular yeast to hyphal transition (YHT) decision making process, we focused on the most studied transcription factors that mediate hyphal initiation (Efg1 and Brg1), hyphal maintenance (Ume6), or inhibit these processes (Nrg1). The model includes four environmental cues known to regulate the YHT, namely pH, farnesol, temperature, and serum. These environmental stimuli regulate the activity of signaling pathways (e.g. cAMP/PKA and ESCRT) that inhibit the expression of *NRG1*, encoding the major YHT transcriptional repressor and/or induce the expression of *EFG1* and *BRG1*, encoding YHT activators. The transcription factors Efg1 and Brg1, in combination with histone acetyltransferases (HATs) and histone deacetylases (HDACs), in turn, induce histone modifications, leading to the activation of downstream HAGs important for the hyphal morphology (e.g., *UME6, HGC1, HWP1, ALS3*, and *ECE1*) [8]. These HAGs encode other transcription factors such as Ume6 that mediate hyphal maintenance, cyclins such as Hgc1 that determine polarized growth at the hyphal tips, and cell wall proteins such as Hwp1, Als3, and Rbt5 that are important for adhesion [7].

The network underlying the model contains several types of nodes, including environmental signals, mRNAs, proteins (signaling proteins, transcription factors, and epigenetic modulators), and processes. These node types are indicated with different symbol shapes and colors in Figure 1. Nrg1 and Efg1 are divided into multiple forms. For Nrg1, the model separately includes the *NRG1* mRNA transcript (NRG1_T), as well as the Nrg1 protein bound to the promoter regions of hyphal-associated genes (Nrg1@HAGs). For Efg1, the model separately includes the *EFG1* transcript (EFG1_T), the Efg1 protein (Efg1), and the Efg1 protein activated as a result of signal transduction (Efg1_active). The latter allows us to encode the negative feedback that active Efg1 has on the transcription of its own gene [9]. We also include three nodes that describe processes: hyphal_initiation, HAG_transcription, and hyphal_maintenance. Activation of the node hyphal_initiation indicates that external signals have impinged on the YHT core network, suppressing yeast-associated nodes, and beginning transcription of hyphal genes. Activation of HAG_transcription indicates that HAGs (e.g., *HGC1* and *HWP1*) are transcribed. Activation of hyphal_maintenance indicates that the cell has entered a state of sustained hyphal growth and elongation as part of a multicellular hypha [7].

**Figure 1.**
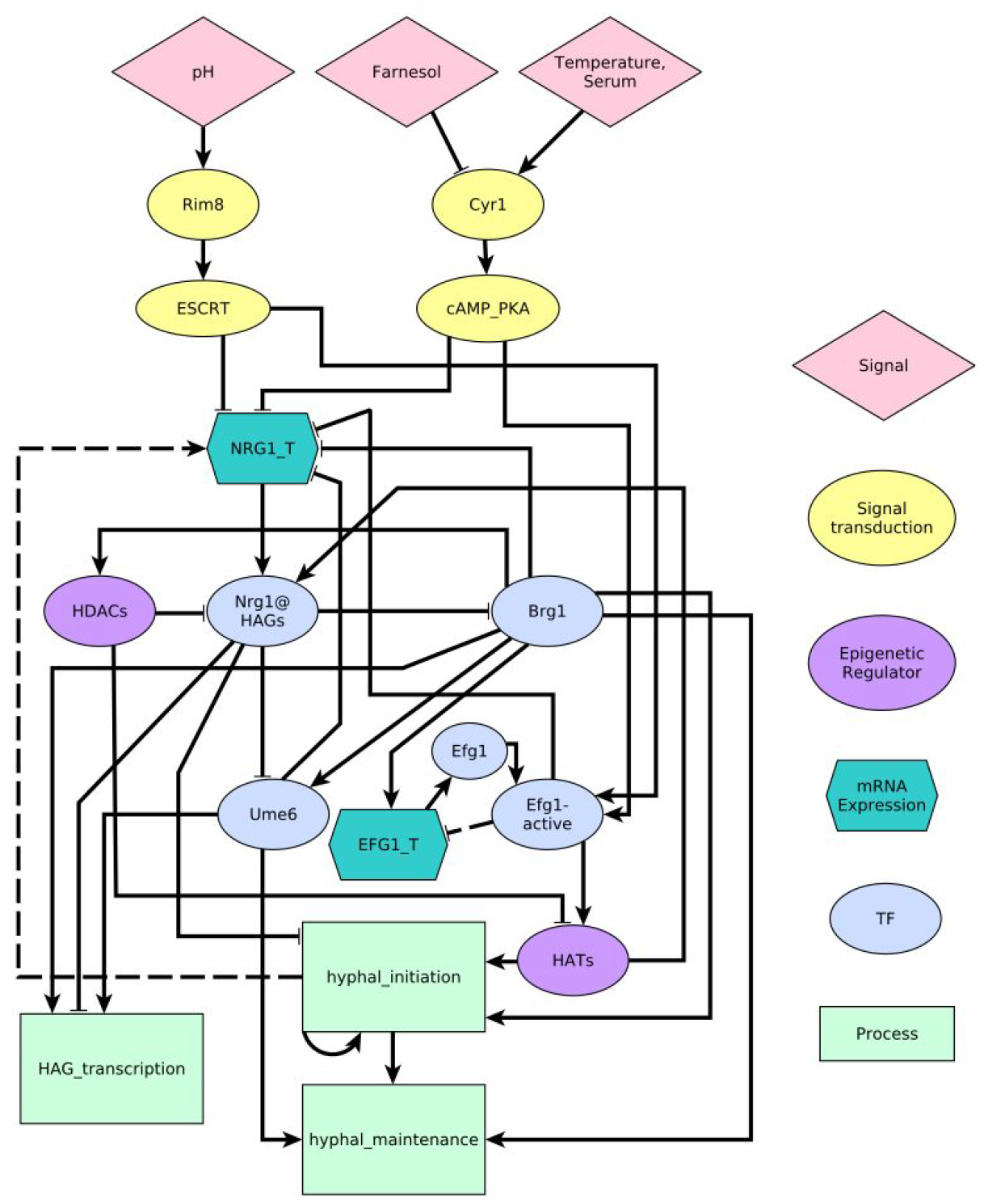
A regulatory network model of the yeast to hyphal transition induced by extracellular signals. The shapes and colors of the nodes indicate their function, as indicated in the key on the right. Dashed edges represent functional relationships whose molecular mechanisms have not been determined. We translate this network into a Boolean dynamic model by characterizing each node with a regulatory function (see Table 1).

To describe the propagation of information in the network from external signals to the ultimate phenotypic output, we formulated a Boolean model. In a Boolean model, each node can be either ON or OFF (1 or 0), and the state of the whole system is given by the state of each node in the network. In general, ON should be interpreted as present, expressed, or active, while OFF indicates the opposite. Special cases include the signal nodes pH and Temperature. The OFF state of the node “pH” indicates an acidic environment (pH < 6), while its ON state indicates an alkaline or neutral environment. Temperature = 0 indicates an environment cooler than 37 °C, while Temperature = 1 indicates an environment at 37 °C.

The regulatory interactions between nodes are given by Boolean regulatory functions describing what the next state of the target node will become based on the current state of its regulators. The regulatory functions were determined from the literature. In cases where detailed knowledge was not available, we generally assumed inhibitory dominant regulatory functions. This means that multiple activating edges are related by the “or” operator, while inhibitory edges are related by the “and not” operator. Specific regulatory functions and evidence in the literature for these functions are summarized in Table 1, and additional notes for each function are provided in Text S1.

**Table 1.**
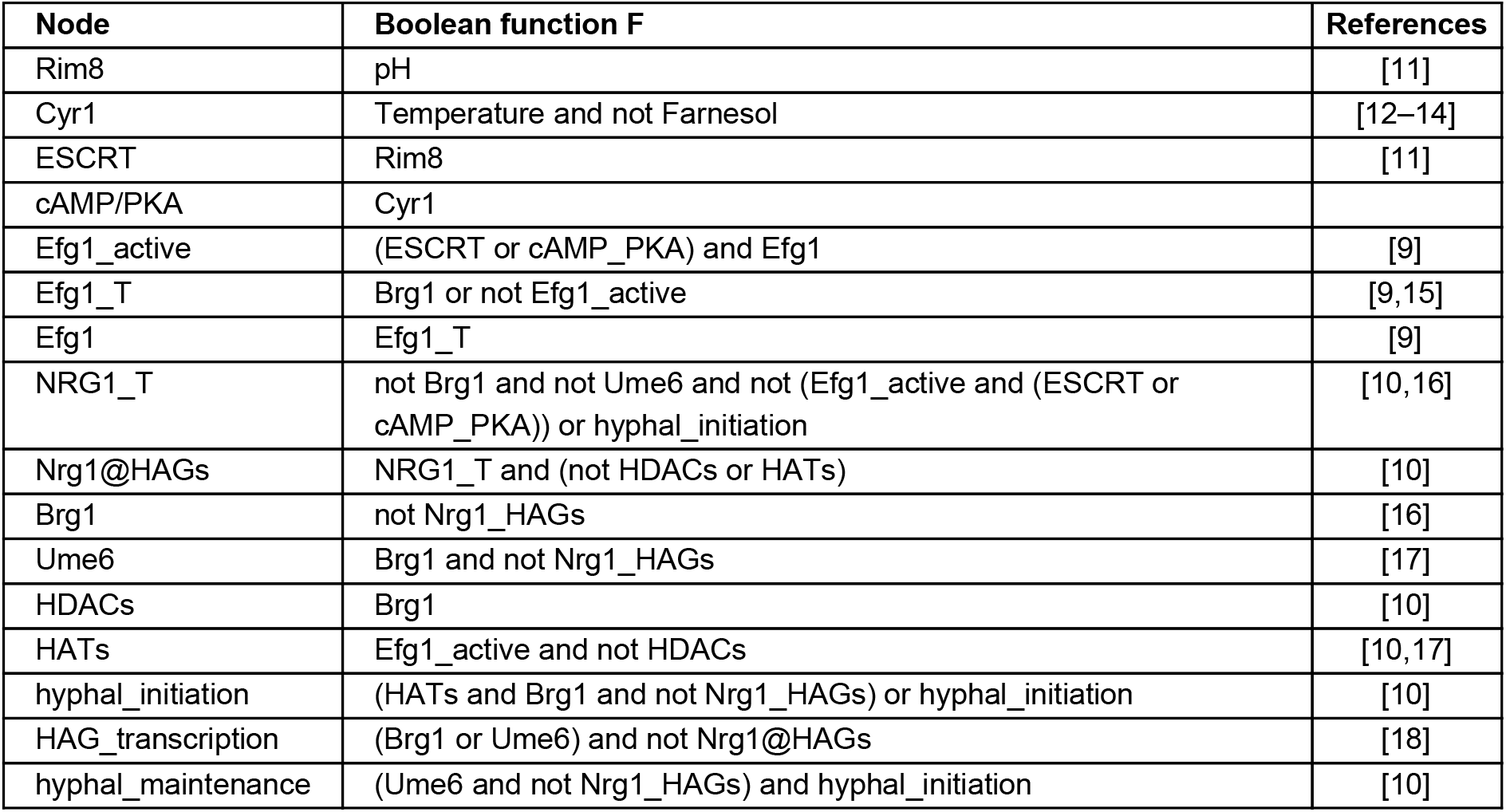
Boolean regulatory functions of each node in the model. Each function indicates the next state of the node as a function of the current state of its regulators. For simplicity the state of each node is represented with the name of the node.

| Node | Boolean function F | References |
| --- | --- | --- |
| Rim8 | pH | [11] |
| Cyr1 | Temperature and not Farnesol | [12–14] |
| ESCRT | Rim8 | [11] |
| cAMP/PKA | Cyr1 |  |
| Efg1_active | (ESCRT or cAMP_PKA) and Efg1 | [9] |
| Efg1_T | Brg1 or not Efg1_active | [9,15] |
| Efg1 | Efg1_T | [9] |
| NRG1_T | not Brg1 and not Ume6 and not (Efg1_active and (ESCRT or cAMP_PKA)) or hyphal_initiation | [10,16] |
| Nrg1@HAGs | NRG1_T and (not HDACs or HATs) | [10] |
| Brg1 | not Nrg1_HAGs | [16] |
| Ume6 | Brg1 and not Nrg1_HAGs | [17] |
| HDACs | Brg1 | [10] |
| HATs | Efg1_active and not HDACs | [10,17] |
| hyphal_initiation | (HATs and Brg1 and not Nrg1_HAGs) or hyphal_initiation | [10] |
| HAG_transcription | (Brg1 or Ume6) and not Nrg1@HAGs | [18] |
| hyphal_maintenance | (Ume6 and not Nrg1_HAGs) and hyphal_initiation | [10] |

Within the scope of the model, serum and temperature have identical downstream effects. For the sake of simplicity, we merged these two environmental signals into a single node, and refer to this node as “Temperature”. Experimental evidence suggests that high temperature and serum are both required to achieve sustained hyphal growth [10]. When comparing the model’s results with experimental findings, we equate the ON state of the input “Temperature” with 37 °C and the presence of serum in the medium.

### The model recapitulates the biological phenotypes and the trajectory of the YHT

We describe the dynamics of the YHT model using two distinct methods: general asynchronous update and stochastic propensity. With both methods, the system evolves until it reaches a stationary state or a group of states that it oscillates within. The term for such final states is “attractor”. Attractors represent stable biological differentiation states or phenotypes. See Methods for more details about the update schemes and attractors.

When considering every combination of states of the three input signals, the YHT network model has 27 attractors (14 if the value of the input signals is not considered), which we broadly categorized as one of four phenotypes: yeast, yeast-like, hyphal-like, and hyphal (Figure 2). Phenotype identification was based on the values of the hyphal_initiation, hyphal_maintenance, and HAG_transcription nodes in the model, as well as on the expression of key transcription factors. One group of attractors corresponds clearly to the yeast state based on the activity of the YHT inhibitor Nrg1 (expressed as the ON state of NRG1_T and Nrg1@HAGs), the inactivity of Brg1 and Ume6, and the lack of hyphal initiation, hyphal maintenance and HAG transcription. The three attractors in this group, marked in blue in Figure 2, only differ in the state of the input nodes Temperature and Farnesol, while the pH must be 0 (acidic environment). We therefore named this group of attractors “yeast”. Another group of attractors (marked by yellow color in Figure 2) has hyphal_initiation = hyphal_maintenance = HAG_transcription = 1. The eight attractors in this group share the activation of hyphal-associated transcription factors and genes and differ only in the state of the signals and of five signal transduction nodes. Therefore, we named this group of attractors “hyphal”.

**Figure 2.**
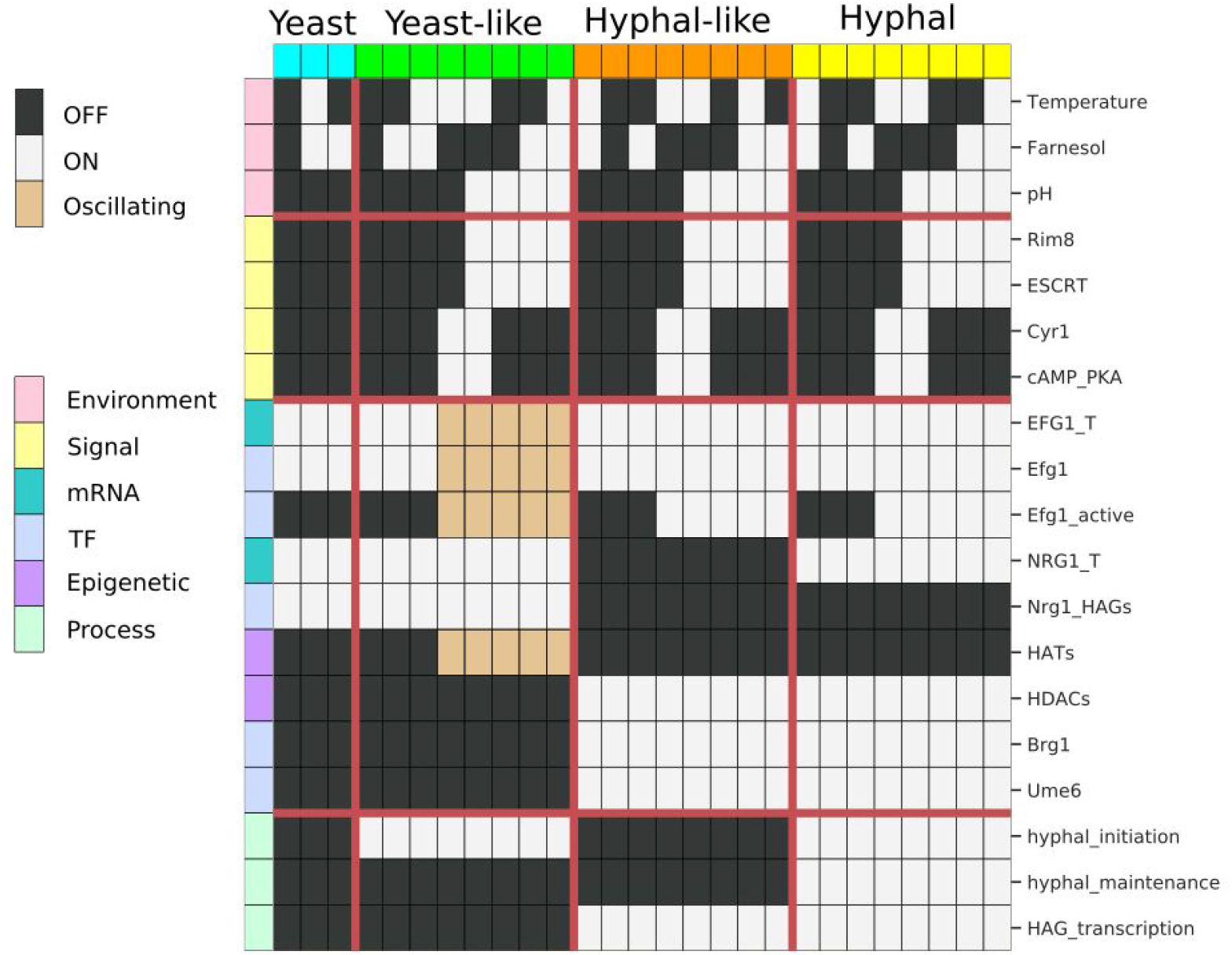
Visual summary of the 27 attractors of the YHT Boolean model. Each row corresponds to a network node and each column indicates an attractor. White indicates active (ON) nodes, black indicates inactive (OFF) nodes, and beige indicates nodes that oscillate. Individual attractors have been assigned to one of four phenotypes − yeast, yeast-like, hyphal-like, and hyphal - based on the status of the hyphal_initiation, hyphal_maintenance, and HAG_transcription nodes (see text).

The other two groups of attractors exhibit intermediate phenotypes. The group of attractors marked in green exhibit active hyphal_initiation, but have active Nrg1 and inactive Brg1 and Ume6, as well as inactive HAG_transcription. These are characteristics of yeast cells, and we therefore named this group of attractors “yeast-like”. A subset of yeast-like attractors exhibit oscillations in Efg1 and HATs, which are driven by the negative feedback loop between Efg1 → Efg1_active -| EFG1_T. We could find no experimental corroboration of this oscillation, although it has been speculated that oscillations caused by this feedback may contribute to variation of *EFG1* expression [19]. Consequently, we do not make any special phenotypic distinction between oscillating and non-oscillating yeast-like attractors.

The group of attractors marked in orange in Figure 2 fail to activate hyphal_initiation and hyphal_maintenance, yet they exhibit expression of *BRG1* and *UME6*, as well as active HAG transcription. Due to the presence of these hyphal characteristics we named this group of attractors “hyphal-like”. These attractors may describe a pseudohyphal phenotype. Unlike hyphae, which only grow at the tip, any cell within pseudohyphae can divide and branch, but, unlike hyphae, the daughter cells remain attached to the mother cells. The formation of pseudohyphae likely involves the same transcriptional core as the YHT, and involves transcription of a subset of HAGs [4]. These features are recapitulated by the hyphal-like attractors of our model.

Depending on the values of the external signals, we found up to four coexisting attractors. For example, when pH = Farnesol = Temperature = 0, there are four possible attractors, which belong to the yeast, yeast-like, hyphal-like and hyphal attractor groups, respectively. In general, the yeast-like, hyphal-like, and hyphal attractor groups contain a stable attractor regardless of the external signals while the stability of the yeast attractor requires pH = 0 and either Farnesol = 1 or Temperature = 0.

To simulate YHT we started in a yeast attractor with pH = Temperature = Farnesol = 0, then set pH = 1. We observed trajectories that converged to any of the three other attractor groups: yeast-like, hyphal-like, and hyphal (Figure 3A-C). A prominent trajectory of our model reproduces the known features of complete YHT in response to alkaline pH: upregulation of *BRG1*, hyphal initiation, HAG transcription, hyphal maintenance (Figure 3C). We verified that setting Temperature = 1 could also induce the YHT, in agreement with the observation that 37 °C induces the YHT [7]. We then undertook a systematic analysis of the outcomes of simulations for every environmental setting using two update schemes (general asynchronous or stochastic propensity). Table 2 indicates the probability of converging into each of the four phenotypes (attractor groups) for every environmental setting when starting from an arbitrary initial state that does not already have hyphal_initiation = 1 or hyphal_maintenance = 1 or from a yeast attractor. In both update schemes, when pH = 0 and either Farnesol = 1 or Temperature = 0 (top row of each table panel) only the yeast and hyphal-like phenotypes are reachable from an arbitrary state, and a system that starts in a yeast state stays in that state. Indeed, yeast is the dominant growth form of C. albicans wild type strains in an acidic environment with temperature lower than 37 °C [4]; in the following we will refer to this environmental condition as a yeast-favoring condition. When either pH = 1, or Farnesol = 0 and Temperature = 1, the yeast attractor is no longer stable, and the system converges into the hyphal-like phenotype, the hyphal phenotype, or to the yeast-like phenotype.

**Table 2.** The probability of converging to each attractor from an arbitrary initial state with hyphal_initiation = 0 and hyphal_maintenance = 0, or from a yeast state. While three attractor groups are stable in any environment, their reachability depends on the environment and on the initial state. For example, in a yeast-favoring environment (first row) if the system starts in a yeast state it will remain in that state. The results are qualitatively the same whether general asynchronous update (top) or stochastic propensity update (bottom) is used.

|  | Initial Condition | Environment | Yeast | Yeast-like | Hyphal-like | Hyphal |
| --- | --- | --- | --- | --- | --- | --- |
| General Asynchronous | Arbitrary* | pH = 0 and (Farnesol = 1 or Temperature = 0) | 31% | 0% | 69% | 0% |
|  |  | pH = 1 or (Farnesol = 0 and Temperature = 1) | 0% | 10% | 62% | 28% |
|  | Yeast | pH = 0 and (Farnesol = 1 or Temperature = 0) | 100% | 0% | 0% | 0% |
|  |  | pH = 1 or (Farnesol = 0 and Temperature = 1) | 0% | 12% | 51% | 37% |
| Stochastic Propensity | Arbitrary* | pH = 0 and (Farnesol = 1 or Temperature = 0) | 26% | 0% | 74% | 0% |
|  |  | pH = 1 or (Farnesol = 0 and Temperature = 1) | 0% | 8% | 54% | 38% |
|  | Yeast | pH = 0 and (Farnesol = 1 or Temperature = 0) | 100% | 0% | 0% | 0% |
|  |  | pH = 1 or (Farnesol = 0 and Temperature = 1) | 0% | 9% | 38% | 53% |

**Fig 3.**
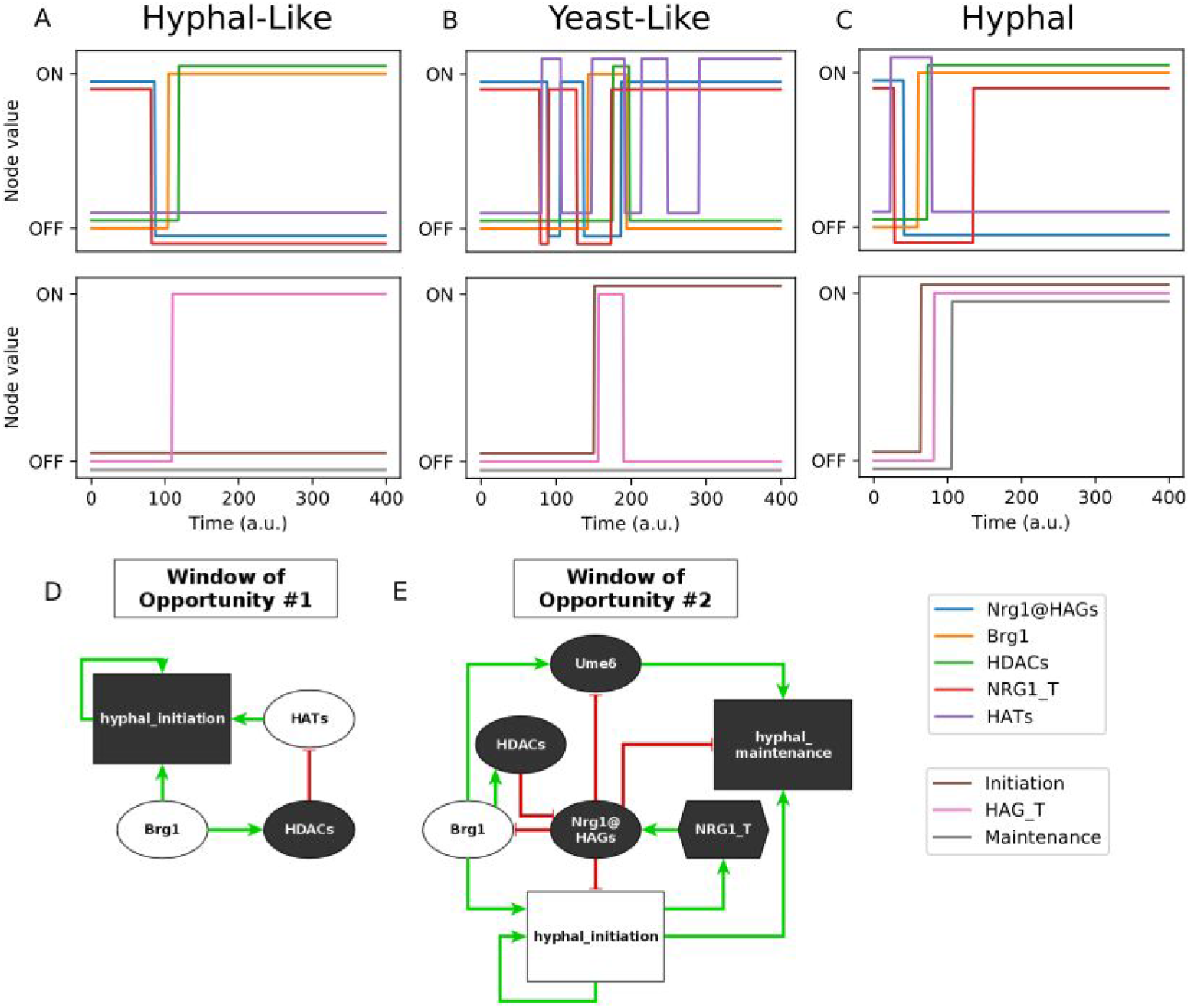
(A-C) Illustrative trajectories of simulated yeast cells placed into an alkaline environment (pH = 1). Simulations use general asynchronous update. (A) A yeast cell fails hyphal initiation, yet hyphal-associated transcription factors (TFs) such as Brg1 are activated and HAGs are transcribed. This trajectory ends in the hyphal-like attractor. (B) A yeast cell achieves hyphal_initiation = 1, and transiently activates HAG_transcription. However the cell fails to lock in hyphal_maintenance, and the yeast program is reestablished. This trajectory ends in the yeast-like attractor. (C) A yeast cell completes the YHT, and ends in the hyphal attractor. (D-E) Motifs from the network controlling these windows of opportunity. White nodes begin as ON at the start of the window of opportunity, while black nodes begin as OFF. (D) Once Brg1 is activated, hyphal_initiation must be activated before HATs are silenced to continue the YHT. If HDACs silence HATs first, the network cannot complete hyphal_initiation, and instead reaches a hyphal-like (pseudohyphal) phenotype. (E) NRG1_T must be silenced to begin hyphal_initiation, but can turn back ON once hyphal_initiation has started. If Nrg1@HAGs reactivates before Brg1 activates HDACs, the system reverts to a yeast-like phenotype. This window may be skipped entirely if HDACs have activated prior to hyphal initiation.

### Stable motif analysis reveals decision points for successful and failed YHT

To understand how the system makes decisions to evolve toward a specific attractor, we performed stable motif analysis [20] on the YHT model (Figure 4). Stable motifs represent subsets of the Boolean network that, once they achieve a certain state, become locked in that state [21]. They are thus the building blocks of attractors. The YHT network has a single unconditional stable motif, which does not depend on environmental conditions. This stable motif consists of hyphal_initiation = 1, which expresses the irreversible nature of hyphal initiation. In addition, there are six conditionally stable motifs. Conditionally stable motifs are only stable motifs if some external condition is met, such as a fixed state of an environmental source node or the stabilization of a parent stable motif. Particularly, the conditionally stable motif outlined in blue involves the activation of the main hyphal inhibitor Nrg1 and the inactivation of the hyphal activators Brg1, Ume6 and HDACs (Figure 4). As can be seen in Figure 1, there is a mutual inhibitory relationship between NRG_T and NRG1@HAGs on one hand, and Brg1, Ume6 and HDACs, on the other hand. This blue conditionally stable motif expresses one of the two possible states of that mutual inhibitory relationship, and is conditioned on the OFF state of both ESCRT and cAMP/PKA, which is true for the three yeast-favoring environmental conditions described by (pH = 0) AND (Farnesol = 1 OR Temperature = 0). The conditionally stable motif hyphal_initiation = 0 can also lock in in the same set of environmental conditions. In a yeast-favoring environment, if the system starts from an initial condition in which hyphal_initiation = 0 (which is the typical case), then this value is stable, and the yeast-like and hyphal phenotypes will be unreachable (this finding is also reflected in Table 2).

**Figure 4:**
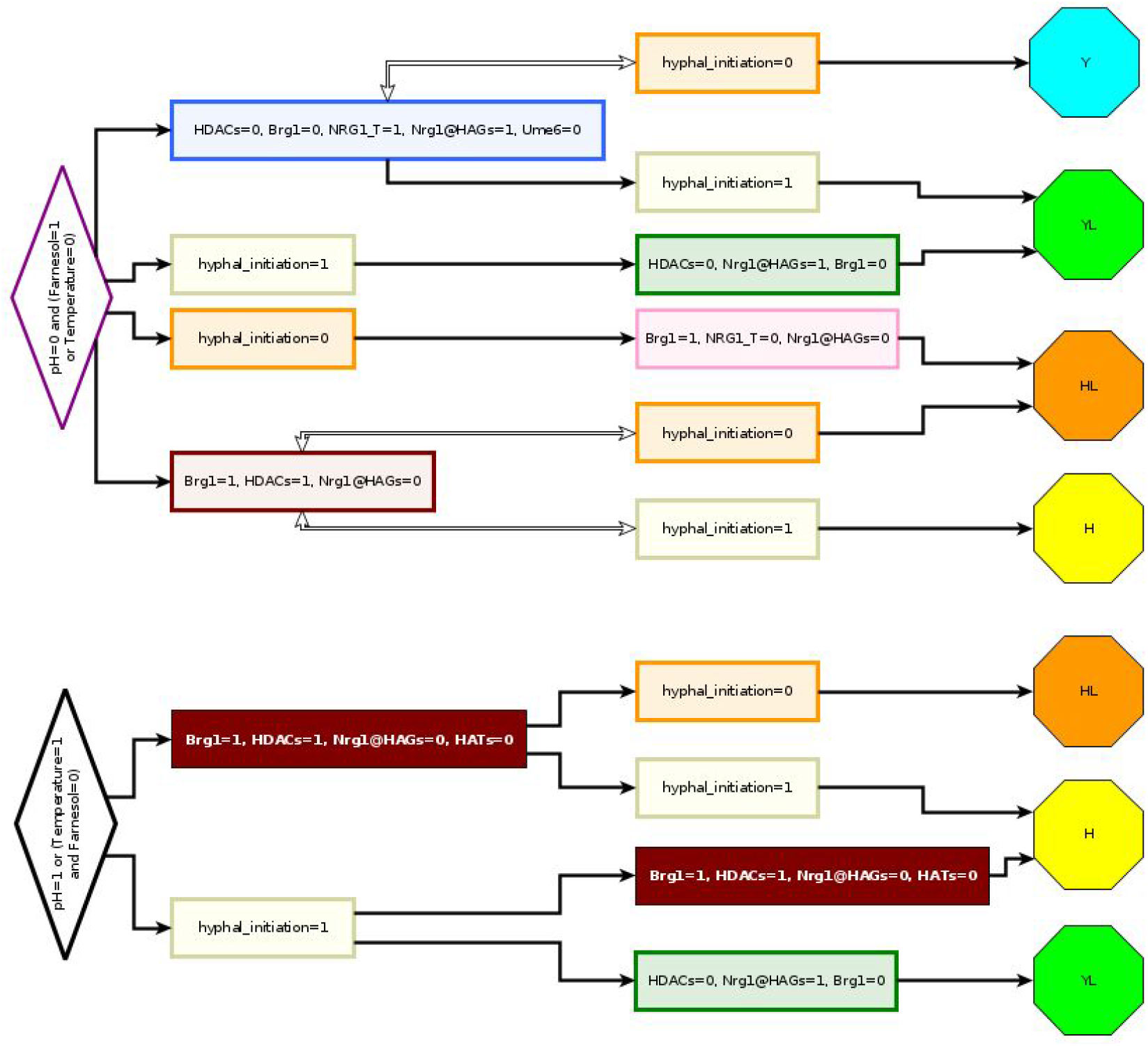
Stable motif succession shows paths to different phenotypes. The colored outlines identify each unique stable motif. From left to right, as subsets of the Boolean model satisfy the shown conditions, they become locked in as stable motifs. For example, in YHT inducing conditions (bottom, such as pH=1), if at any point the Brg1 = HDACs = 1, while simultaneously Nrg1@HAGs = HATs = 0, then the dark brown conditionally stable motif becomes locked in. Further, once it is locked in, either of the conditionally stable motifs hyphal_initiation = 0, or hyphal_initiation = 1, may become locked in. Depending on which path is taken, the system will then evolve toward either the hyphal attractor (if hyphal_initiation = 1 locks in) or the hyphal-like attractor (if hyphal_initiation = 0 locks in).

A conditionally stable motif expressing the activity of Brg1 and HDACs and the inactivity of the hyphal inhibitor Nrg1@HAGs has two variants. The first variant, shown in brown outline, is a stable motif in the environmental conditions (pH = 0) and (Farnesol = 1 or Temperature = 0). The second variant, shown with a brown background, also includes the inactivity of HATs. This variant is a stable motif in the environmental conditions (pH = 1) or (Farnesol = 0 and Temperature = 1). The conditionally stable motif outlined in green is a subset of the blue conditionally stable motif and is conditioned on hyphal_initiation = 1. The conditionally stable motif outlined in pink overlaps with the brown stable motif, and is conditioned on hyphal_initiation = 0.

The sequence of which subsequent stable motifs may lock in after a given stable motif locks in is shown in the stable motif succession diagram [21] (Figure 4). The succession diagram confirms the simulation results that the yeast attractor is only reachable when pH = 0, and either Farnesol = 1 or Temperature = 0. The trajectory toward the yeast attractor involves the stabilization of the blue conditionally stable motif and the hyphal_initiation = 0 conditionally stable motif. These motifs are independent of each other and thus could activate in either order in an arbitrary trajectory; this is indicated by the bidirectional arrow in Figure 4. The other three attractors are also reachable in these environmental conditions through the successive lock-in of two conditionally stable motifs. The reason that the simulations reported in Table 2 only reach the yeast or hyphal-like attractors is that those simulations have hyphal_initiation = 0 in the initial condition, which is a stable motif under these environmental conditions, and thus is immediately locked in, restricting the allowed successions.

In the hyphal-inducing environmental conditions (pH = 1 or Farnesol = 0 and Temperature = 1) the locking in of the brown conditionally stable motif can be paired with either state of hyphal_initiation. According to the regulatory function for hyphal_initiation (Table 1), locking in the brown stable motif takes away the possibility of hyphal_initiation to turn ON if it was initially OFF. If hyphal_initiation turns on prior to the locking-in of the brown stable motif, the system converges into the hyphal phenotype. As soon as the brown stable motif locks in, hyphal initiation is prevented from turning ON, and thus it will lock in the OFF state; the system will converge to the hyphal-like attractor. The locking-in of hyphal_initiation=1 can be followed up by the brown stable motif or the green conditionally stable motif, which expresses the state opposite of the brown stable motif. The first succession leads to the hyphal phenotype, while the second leads to the yeast-like phenotype.

### Stable motif decision points correspond to YHT windows of opportunity

In the stable motif succession diagram, when there are multiple edges emerging from a single motif, the system taking one of these edges may represent an irreversible commitment to one of two mutually exclusive trajectories. Figure 4 shows two branch points that determine commitment relevant to the YHT: the edges emerging from the solid brown conditionally stable motif to either the tan or orange motifs indicate commitment to either the hyphal or hyphal-like attractor groups. Similarly, the edges emerging from the tan stable motif to either the solid brown or green motif dictate commitment to the hyphal or yeast-like attractor groups. The choice of one path versus another depends on the timing of specific events.

The first branch point depends on a sequence of events starting when Brg1 turns ON. While Brg1 remains ON, deactivation of HATs will follow via the inhibitory path Brg1 → HDACs -| HATs. However, activity of HATs is a requirement for hyphal_initiation to turn ON. If hyphal_initiation activates before the node HATs turns OFF, then the system will follow the path toward the hyphal attractor group. Conversely, if HATs turns OFF before hyphal_initiation activates, the system will proceed toward the hyphal-like attractor group. For example, in the trajectory in Figure 3A, HATs turn OFF before hyphal_initiation turns ON, causing the system to proceed to the hyphal-like phenotype. This corresponds to the small incoherent feedforward loop illustrated in Figure 3D.

The second branch point depends on the timing of events following the activation of hyphal_initiation. If the brown motif (which contains Nrg1@HAGs = 0) locks in, or has locked in prior to the activation of hyphal_initiation, then the system will proceed to the hyphal attractor group. Yet, NRG1_T returns once hyphal_initiation activates, and can lead to activation of Nrg1@HAGs, which is part of the green motif. Nrg1@HAGs activity is sufficient to lock in the green motif, which expresses the deactivation of the core hyphal program, leading to the yeast-like attractor group. Thus the YHT depends on a race to exclude Nrg1 from the promoter region of HAGs following the reactivation of *NRG1* transcription. This race corresponds to multiple negative regulatory pathways between hyphal_initiation and hyphal_maintenance (Figure 3E), mediated through Nrg1@HAGs.

These decision points, and their corresponding races, reflect the documented *C. albicans* YHT “window of opportunity” [7,10]. This is a transient period, beginning with the downregulation of *NRG1* and ending with the subsequent re-expression of *NRG1*, in which the hyphal program may be established. If it does not establish prior to the window closing, the cells do not complete the YHT. As described above, our model reproduces this behavior, resolving the window of opportunity into two distinct decision points (one regarding hyphal initiation and the other regarding hyphal maintenance), and describing the specific mechanisms by which the window can be missed.

### Network control predictions

Using the Boolean YHT model, we sought to identify interventions, such as controlling the state of one or more nodes or deleting or activating an edge, that could prevent the YHT. Interventions were identified using several different control strategies and objectives, summarized in Table 3. We have applied feedback vertex set (FVS) control, stable motif control, and algebraic edge control with canalizing function analysis to our network model, as well as systematic simulations of perturbations.

**Table 3.** An overview of the control prediction approaches we apply to the *C. albicans* YHT network, their objectives, and the types of interventions they require.

| Control Method | Objective | Interventions |
| --- | --- | --- |
| FVS Control [22] | Force system into target pre-existing attractor | Permanently control source nodes, temporarily control other nodes |
| SM Control [20] | Force system into target pre-existing attractor | Temporarily (and sequentially) control nodes |
| Simulation | Block YHT | Permanently control single node |
| Algebraic Edge Control [23] | Block YHT | Permanently control single edge |

### Feedback vertex set control

A feedback vertex set (FVS) is a collection of nodes in a network whose removal results in a network with no cycles (no feedback loops). On a network with no feedback loops, dynamical processes described by Boolean or differential equation models have a single attractor [22,24]. FVS control thus predicts that by fixing all nodes in a given FVS, as well as all source nodes, to match a particular attractor, one can force the system from any state into that attractor [25]. Once the system achieves that target attractor, control of the FVS nodes may be relaxed, though control of the source nodes must be maintained. Unlike the other control methods, FVS only requires knowledge of the network’s topology (Figure 1), that is, the collection of nodes and edges, as well as knowledge of the attractors, but it otherwise requires no specific details of the regulatory functions.

The YHT network contains a strongly connected component (feedback-rich subgraph) of 10 nodes. The FVS of the network consists of three nodes: Nrg1@HAGs, at least one node from the Efg1 feedback loop, and hyphal_initiation (which has a self-loop). FVS control predicts that a control strategy for ensuring that the system converges into the yeast attractor is to maintain Nrg1@HAGs = 1, Efg1 = 1, and hyphal_initiation = 0 to eliminate feedback sets, as well as ensure pH = 0 and either Farnesol = 1 or Temperature = 0 to control source nodes (see panel A of Figure S1). Conversely, FVS control into the hyphal attractor group requires setting Nrg1@HAGs = 0, Efg1 = 1, and hyphal_initiation = 1 (panel B of Figure S1). As this attractor group is reachable in any environmental condition, the source nodes do not need to be controlled.

FVS control provides a sufficient condition to maintain a given attractor. Nevertheless, it may be that a subset of nodes can still accomplish the control objective. This is especially important to identify here, as fixing nodes such as hyphal_initiation may have no obvious biological implementation.

### Stable motif based control

Stable motif control seeks to determine a sequence of driver nodes that, if transiently maintained in a fixed state, will lock in a sequence of stable motifs that will force the system into a desired attractor from any initial condition. A variation of stable motif control also identifies sequential control of driver nodes that drives the system into one of multiple target attractors [26].

Figure 5 shows driver sequences needed to drive the system into an attractor with hyphal_maintenance = HAG_transcription = 0 (corresponding to one of the yeast or yeast-like attractor groups), or hyphal_maintenance = 1 (corresponding to a hyphal phenotype). Unlike FVS control, stable motif control is able to force the system to specific attractor groups by only controlling one or two nodes, depending on the control objective and environmental conditions. For example, temporarily controlling hyphal_initiation = 1 (after which it locks in), followed by temporarily controlling Brg1 = 0 (until the green stable motif in Figure 4 locks in) is sufficient to achieve hyphal_maintenance = HAG_transcription = 0 in any environment. In this case, the system will follow a trajectory toward a yeast or yeast-like attractor. Driving the system to a hyphal phenotype in any environment and from any initial condition requires temporarily holding hyphal_initiation = 1 (after which it locks in), followed by holding any node value of the brown-outlined motif (NRG1@HAGs = 0, HDACs = 1, or Brg1 = 1). The stable motif based control sets still have the shortcoming of involving direct control of hyphal initiation, which is difficult to implement experimentally.

**Figure 5.**
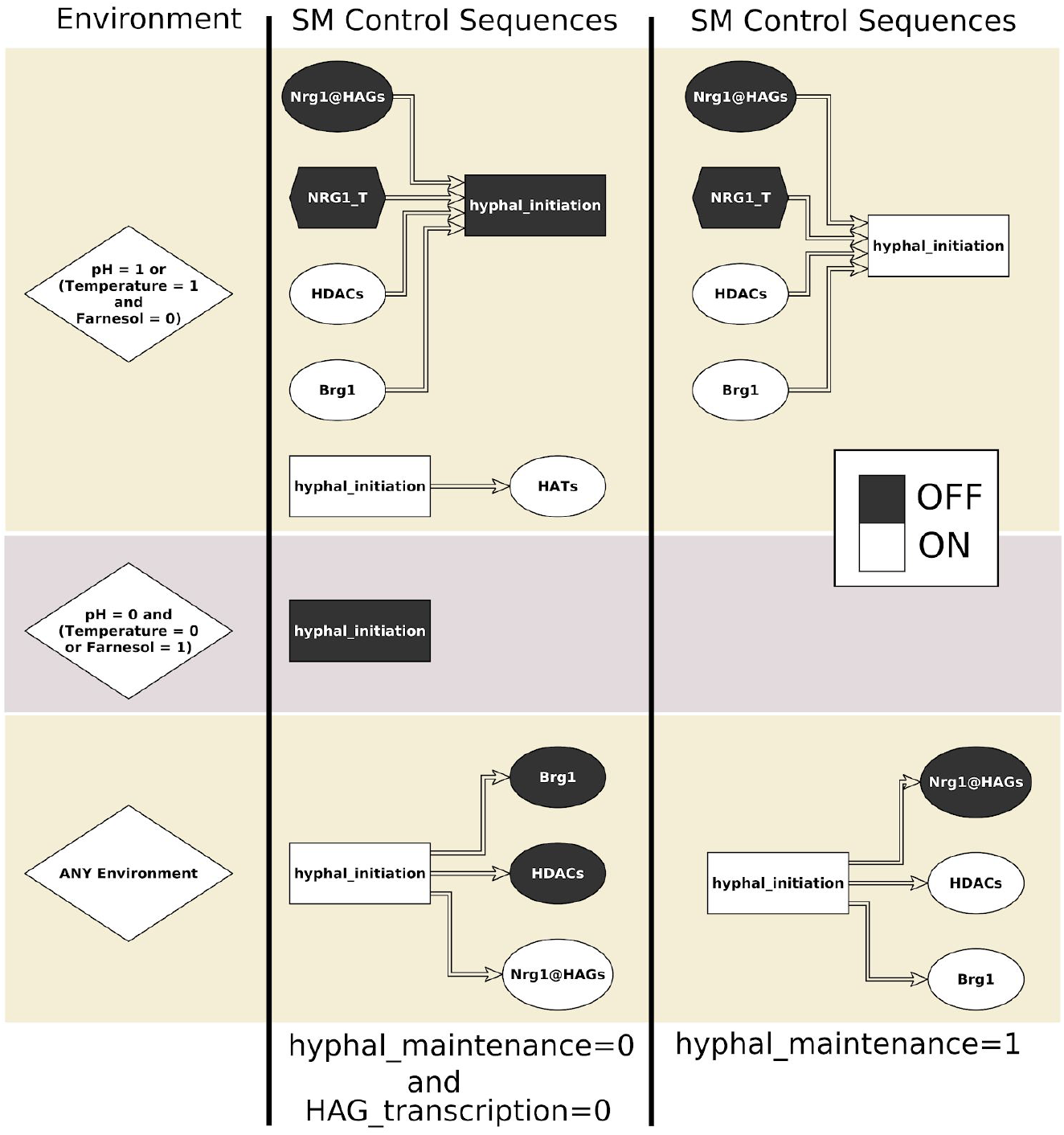
Stable motif control strategies for different environmental conditions and control objectives. Controls are implemented from left to right, where each control should be maintained until the corresponding motif locks in, then the next applied, and so on. When there are multiple different control options, any single one is sufficient.

### Simulated perturbations

We systematically simulated permanent deletions (holding in the state 0) and activations (holding in the state 1) of the nine nodes that are not signals, signaling intermediaries, or phenotypic outcomes using both general asynchronous and stochastic propensity updating schemes. To identify interventions that block YHT, we began the simulations in a yeast state placed into an environment with Farnesol = Temperature = 0, pH = 1. As indicated in Table 2, the unperturbed system undergoes the YHT in about 37% of trajectories for general asynchronous update (53% for stochastic propensity update), otherwise missing one of the two windows of opportunity. We found 8 node interventions that ensure none of the trajectories converge to a hyphal phenotype (Table 4). Among these, the interventions Brg1 = 0, HDACs = 0, Ume6 = 0, or Nrg1@HAGs = 1 led to the complete elimination of the hyphal attractor. Indeed, it was found experimentally that deletion of *BRG1* [27], *HDA1* [10], or *UME6* [28] led to impairment of the YHT. These four states are incompatible with both variants of the brown motif, whose locking-in is necessary for the hyphal attractor. Instead, the trajectories starting from the yeast attractor converge into either a hyphal-like or yeast-like attractor (in case of Ume6 = 0) or solely to a yeast-like attractor (in the other three cases). In the remaining four cases the hyphal attractor exists but it is not reachable from an initial condition corresponding to yeast. In the case of deletion of *EFG1*, the system stays in the yeast attractor, and for HATs = 0 it converges into a hyphal-like attractor. Indeed, it was found experimentally that deletion of *EFG1* prevented hyphal formation [29] and deletion of *YNG2*, encoding the Yng2 subunit of the HAT NuA4 complex led to diminished HAG transcription and significantly impaired hyphal formation [17].

**Table 4.**
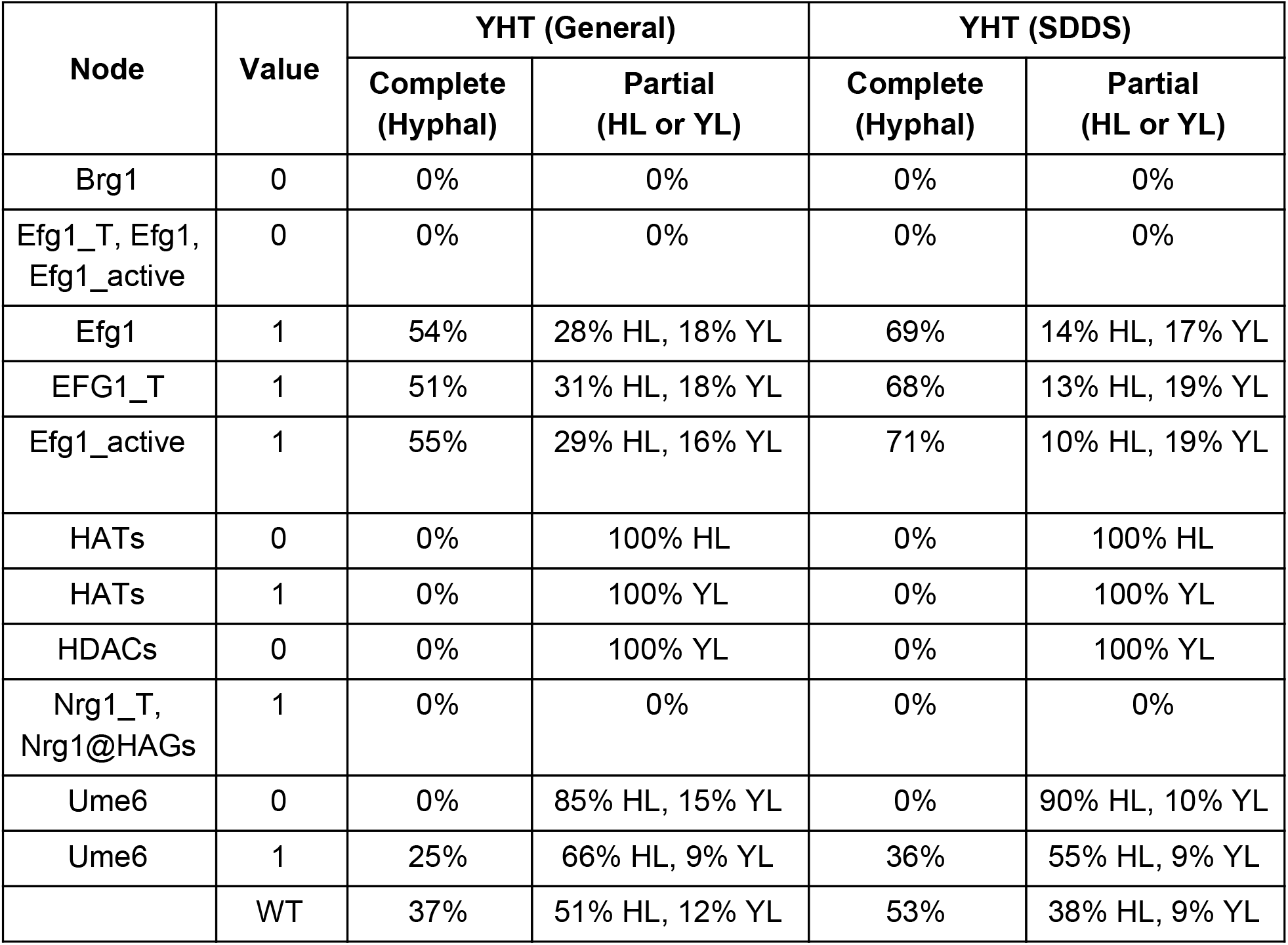
Probabilities of completing the YHT in the condition pH = 1 and Farnesol = Temperature = 0, under single-node interventions or WT. Transition probabilities are estimated from 500 simulated trajectories starting from the yeast attractor corresponding to the given environment (see Figure 2). When a perturbed system’s attractor is different from the WT attractor, it is classified into a phenotype (attractor group) by similar criteria as in Figure 2; see Figure S2 for more details.

| Node | Value | YHT (General) |  | YHT (SDDS) |  |
| --- | --- | --- | --- | --- | --- |
|  |  | Complete (Hyphal) | Partial (HL or YL) | Complete (Hyphal) | Partial (HL or YL) |
| Brg1 | 0 | 0% | 0% | 0% | 0% |
| Efg1_T, Efg1, Efg1_active | 0 | 0% | 0% | 0% | 0% |
| Efg1 | 1 | 54% | 28% HL, 18% YL | 69% | 14% HL, 17% YL |
| EFG1_T | 1 | 51% | 31% HL, 18% YL | 68% | 13% HL, 19% YL |
| Efg1_active | 1 | 55% | 29% HL, 16% YL | 71% | 10% HL, 19% YL |
| HATs | 0 | 0% | 100% HL | 0% | 100% HL |
| HATs | 1 | 0% | 100% YL | 0% | 100% YL |
| HDACs | 0 | 0% | 100% YL | 0% | 100% YL |
| Nrg1_T, Nrg1@HAGs | 1 | 0% | 0% | 0% | 0% |
| Ume6 | 0 | 0% | 85% HL, 15% YL | 0% | 90% HL, 10% YL |
| Ume6 | 1 | 25% | 66% HL, 9% YL | 36% | 55% HL, 9% YL |
|  | WT | 37% | 51% HL, 12% YL | 53% | 38% HL, 9% YL |

In the case of simulated constitutive activation of Ume6 the YHT propensity decreased compared to wildtype. This happens in the model because Ume6 inhibits Nrg1@HAGs, decreasing the YHT window of opportunity. In contrast, simulated constitutive activation of the *EFG1* transcript or Efg1 protein increased the YHT propensity. This result is consistent with the hyphal morphologies observed when *EFG1 is* overexpressed [30].

Unlike the previous control methods, which by design force the system into a pre-existing attractor, this method can introduce new attractors. For example, a simulated deletion of *BRG1* prevented the YHT, consistent with experimental observations of defective hyphal elongation in mutants lacking *BRG1 [27]*, and introduced a new yeast-like attractor in which NRG1_T and Nrg1@HAGs oscillate. The new attractors observed for permanent deletions and activations of single nodes are indicated in Figure S2.

We also identified four perturbations that eliminate the yeast attractor even in yeast-favoring environmental conditions, namely Brg1 = 1, HDACs = 1, NRG1_T = 0, and Nrg1@HAGs = 0. These states are incompatible with the blue motif (Figure 4), whose locking in is necessary for the yeast attractor. Any of these node states can ensure the locking in of both versions of the brown conditionally stable motif in Figure 4. When this motif locks in, the current state of hyphal_initiation determines whether the system converges to the hyphal-like or hyphal attractor groups. Hyphal initiation following a change in environment requires that hyphal-inducing signals propagate through two parallel pathways: activating the brown motif (via NRG1_T downregulation), and activating HATs (via Efg1). Due to the proximity of the controls to the brown motif, in all simulations the brown motif locked in before HATs turned on, causing the system to converge to a hyphal-like attractor. Thus, our model predicts that these perturbations would lead to filamentation (likely pseudohyphae) even in yeast-favoring environmental conditions. Indeed, experiments indicate that deletion of *NRG1* leads to pseudohyphae [31,32]. In contrast to the model prediction, *C. albicans* cells engineered to ectopically express *BRG1* under yeast-favoring conditions stayed in a yeast phenotype [27]. One potential explanation of this discrepancy is that Brg1 translated from ectopically expressed *BRG1* may not be able to recruit HDACs to the promoter region of HAGs to exclude Nrg1@HAGs; indeed the same study found that in strains ectopically expressing *BRG1*, the Brg1 protein could not bind to the promoter region of HAGs.

### Algebraic Control with Edge Knockouts

Algebraic control [33] uses the polynomial form of the Boolean functions. Two types of control objectives can be formulated: node control and edge control. The identification of control targets is achieved by encoding the nodes of interest as control variables within the functions; edges of interest are encoded as control variables within the inputs of the functions. Then, the control objective is expressed as a system of polynomial equations that is solved by computational algebra techniques. For node control, we considered the environmental condition pH = Rim8 = 1, Temperature = Farnesol = Cyr1 = cAMP_PKA = 0 and set the objective of finding node knockouts or constitutive activations for which there is an attractor of the system that has the hyphal_maintenance and HAG_T nodes OFF. Thus, we set our objective to find controls such that 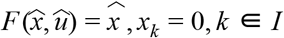, where the index set *I* corresponds to the indexes of hyphal_maintenance and HAG_T. We found the following node controls: Brg1 = 0, HDACs=0, and NRG1@HAGs=1. These interventions were also identified by simulations to block the YHT.

For edge control, we set the control objective of destroying (or blocking) the fixed point *x* _0_ corresponding to the hyphal state. Thus, our goal is to find controls *u* such that *F* (*x* _0_, *u*) =/ *x* _0_. Using the edge control approach we identified the following edge deletions and activations, shown in Table 5, that are effective for blocking transition into the hyphal state.

**Table 5.** Edge perturbations predicted by algebraic control to block the YHT. Total effect measures the percent change in the state transition graph, and time to absorption (TTA) measures how long it takes to reach an attractor, starting from the yeast state, setting pH=1.

| Source | Target | Control type | Description | Total Effect | TTA (WT = 16.8 steps) |
| --- | --- | --- | --- | --- | --- |
| Brg1 | Ume6 | Deletion | Equivalent to deletion of Ume6 | 12.5% | 16.4 |
| Nrg1@HAGS | Ume6 | Activation |  |  |  |
| HATs | Nrg1@HAGS | Activation | Blocks hyphal attractor with 75% YL, 25% HL. | 6.25% | 18.71 |
| HDACs | Nrg1@HAGS | Deletion |  |  |  |
| Nrg1@HAGS | Brg1 | Activation | Equivalent to deletion of Brg1; makes the yeast state an attractor | 25% | 0 |
| Brg1 | HDACs | Deletion | Equivalent to deletion of HDACs | 25% | 21.34 |
| HDACs | HATs | Deletion | Blocks hyphal attractor with 100% YL | 12.5% | 24.12 |

For each edge control, we calculated the total effect (the total change in the state transition graph) as described in the Methods. Interventions with larger total effect induce greater systemic changes in the state transition graph of the system, and therefore may be associated with more side effects [23]. We likewise calculated the system’s time to absorption (TTA), which corresponds to how long it takes the system to reach an attractor, starting from the yeast state, and setting pH = 1. Controls that have a lower TTA indicate that the system will quickly reach the attractor. In the case when TTA=0, the perturbed system has a stable yeast attractor even when pH = 1. This is the case for constitutive activation of the inhibitory edge Nrg1@HAGs-| Brg1.

The edge interventions with smallest total effect (6.25%) are constitutive activation of the HATs → Nrg1@HAGs edge, or constitutive deletion of the HDACs -| Nrg1@HAGs edge. The effect of these interventions ensures that if Nrg1 is expressed, it will bind to HAGs, thus under this intervention the system misses the second window of opportunity (Figure 3E). With this intervention, the system converges preferentially into the YL attractor, or alternatively the HL attractor, with a slightly longer convergence time than the WT system would converge to a hyphal attractor.

### Experimental verification of predicted control interventions

We performed two new experiments to test two model-predicted interventions that eliminate the YHT: deletion of *UME6* and deletion of the HDAC *HDA1*. The model predicts both of these interventions make hyphal maintenance impossible, though it predicts differences in the final attractors. Setting Ume6 = 0 leads to a three-attractor repertoire: yeast, yeast-like, and a novel attractor group named hyphal-like 2 (see Figure S2). The attractors of the Ume6 = 0 system do not achieve hyphal maintenance but do exhibit one or both of the other hyphal-associated phenotypic outcomes. Setting HDACs = 0 in the model leads to convergence to a yeast-like attractor, which is a much stronger departure from the unperturbed system’s outcome. We determined the morphology of cells of *C. albicans* strains lacking *UME6* and *HDA1* at 90 minutes after inoculation. By this time, cells of the wildtype *C. albicans* strain have completed hyphal initiation and are in the elongation phase [7]. The experimental condition was neutral (pH = 7.0) RPMI-1640 medium at 37 °C; these mutant strains were not tested under these conditions in any prior publications we found. As shown in Figure 6, compared to wildtype strain at 90 minutes (0% yeast-form, 0% transitional-form, 100% true hyphal-form), the *ume6* Δ/Δ strain displayed a relatively minor filamentation defect (4% yeast-form, 29% transitional-form, 67% true hyphal-form), while the *hda1* Δ/Δ strain displayed a severe filamentation defect (60% yeast-form, 40% transitional-form, 0% true hyphal-form). These results are consistent with the model predictions, especially in terms of the relative severity of the two defects.

**Figure 6.**
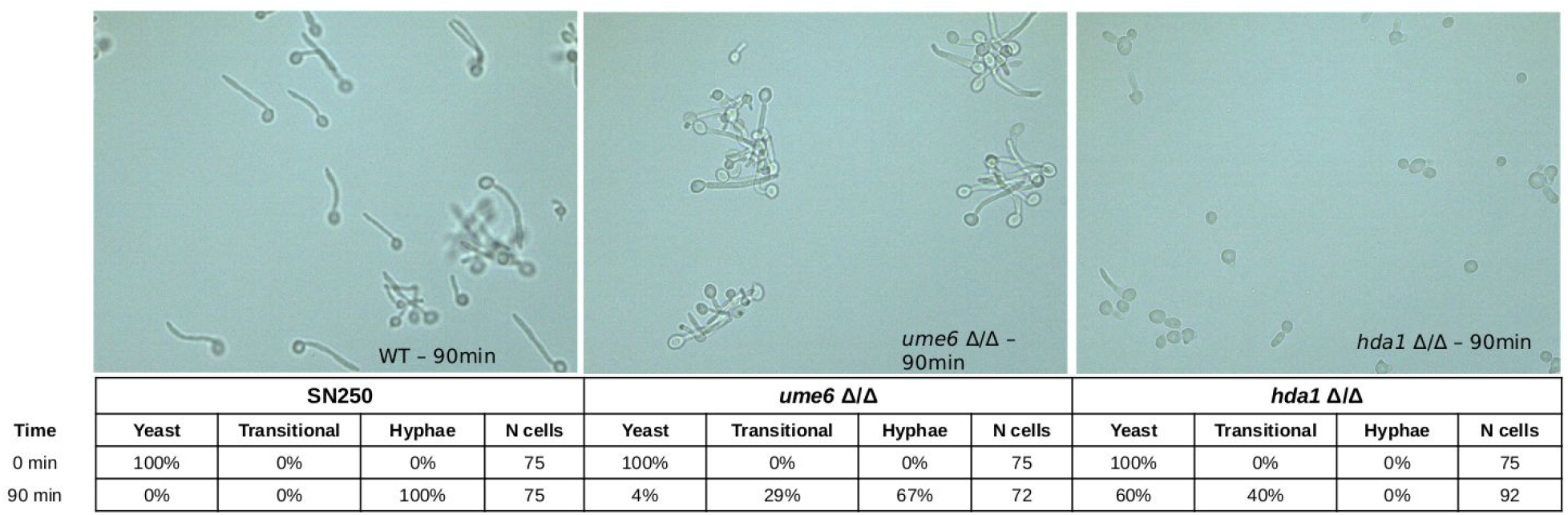
Representative images and quantification of the experimentally observed growth phenotypes of the wildtype strain SN250 (left), *ume6* Δ/Δ strain (middle), and *hda1* Δ/Δ strain (right) *C. albicans* strains grown in RPMI medium at 37 °C, pH 7.0.

As an additional verification of the model, we compared model-predicted outcomes of simulated controls with published results of corresponding experimental interventions. As shown in Supplemental Table 1, the model and experiments agree in 18 out of 22 cases. The four cases of disagreement pertain to *NRG1* deletion and *BRG1* deletion under filamentation-inducing conditions, and Efg1 constitutive activation and Brg1 constitutive activation under conditions favoring the yeast-form.

The model predicts that deletion of *NRG1* leads predominately to a hyphal-like attractor in every environment, while experimentally it was found that in hyphal-inducing conditions, deletion of *NRG1* led to hyphal formation [31]. In the model, deletion of *NRG1* often leads to early activation of HDACs, which precludes the temporary activation of HATs, which in turn is necessary for hyphal initiation (Figure 3D). This discrepancy could be mitigated by revising certain assumptions of the model. Activation of Brg1 may require the action of a specific activator in addition to Nrg1@HAGs inactivity; indeed Brg1 is documented to be part of a network of six cross-activating transcription factors [29], which also includes Efg1. Further analysis of the functional requirement for this network and its relationship with Nrg1 would elucidate the regulatory function of Brg1. Another possibility is that the requirement for HATs activity for hyphal initiation may be less strong than assumed in the model.

Constitutive activation of Efg1 (simulated by maintaining Efg1_active = 1) in a yeast-favoring environment leads to an attractor that exhibits a low level of Efg1_T and Efg1 but otherwise is the same as the unperturbed system’s yeast attractor. In contrast, a pseudohyphal phenotype was observed experimentally [30]. The reason the model does not recapitulate this result is the assumption that Nrg1_T downregulation requires active Efg1 in collaboration with cAMP/PKA or pH signaling. The discrepancy would be resolved by the alternative assumption that the active Efg1 is the mediator of the effect of the environment on Nrg1_T. Indeed, a model version that omits the direct effects of the environment on Nrg1_T indicates a mixture of hyphal and hyphal-like attractors in a yeast-favoring environment.

The simulated deletion of Brg1 leads to the system staying in a yeast state in hyphal-inducing conditions, while experimentally it was found that a *BRG1* deletion strain exhibited competent germ tube formation and defective hyphal elongation. The discrepancy stems from the assumption that Brg1 activity is required for hyphal initiation. It is possible that other transcription factor(s) within the six transcription factor network mentioned above [29] could rescue hyphal initiation in the absence of Brg1.

Constitutive activation of Brg1 in a yeast-favoring environment leads to a hyphal-like attractor in the model while experimentally overexpressing *BRG1* caused the cells to remain in the yeast phenotype, perhaps because the ectopically induced Brg1 was unable to bind to the promoters of HAGs [27]. The discrepancy could be mitigated by a condition for the activation of HDACs that requires more than the presence of Brg1.

## Discussion

Here we have developed a new Boolean model of the *C. albicans* YHT, and demonstrated that it recapitulates several known behaviors. Our model has attractors corresponding to the yeast and hyphal phenotypes, as well as two types of attractors that correspond to different ways the system can fail to complete the YHT, by failing to pass so-called “windows of opportunity” (Figure 2). The hyphal-like attractors exhibit expression of *BRG1, UME6* and hyphal-associated genes, but fail to activate hyphal initiation. Our model predicts that the YHT can arrest if histone acetyltransferases (HATs) inactivate prior to the transcription of hyphal-associated genes. This state is a point attractor in our model, which may correspond to a pseudohyphal phenotype, or it may not be a stable biological state. Further experimental investigation of this system may support or contradict the stability of this state. It may also reveal the conditions under which the transition can resume or alternatively the system can reset to a yeast state. The yeast-like attractors exhibit active hyphal initiation, but have active Nrg1 and inactive Brg1 and Ume6, which are characteristic of yeast cells. We interpret these states as the YHT arrested after hyphal initiation, followed by a resetting into the yeast-like state of the transcriptional regulators.

Through stable motif analysis, we showed that the previously described YHT window of opportunity corresponds to two branch points in stable motif succession (Figure 4). The choice of which branch the system takes depends on the specific timing of events within two small subnetworks we identified as the window of opportunity motifs (Figure 3). Both of these subnetworks contain an incoherent feed-forward loop, which is the coexistence of two short paths of opposite signs between a pair of nodes. The timing of the events in each path determines the outcome. In the first window of opportunity (Figure 4D), hyphal_initiation must be activated before HATs are silenced by HDACs to continue the YHT. In the second window of opportunity (Figure 4E), HDACs must be activated by the time the Nrg1_T expression starts to increase again, to avoid the reactivation of Nrg1@HAGs, otherwise the YHT cannot be sustained. Further experimental investigation of these epigenetic regulatory processes will be able to determine their timing and regulation. For example, one may elucidate the timing requirement of the HAT activity by engineering a a timable deletion of *YNG2* (encoding the active subunit of the HAT NuA4) or by including an inducible promoter for a constitutively acetylated (and thus non-responsive to HDACs) mutant (K175Q) *YNG2* in a WT *YNG2* strain.

We applied four methods to predict control strategies of the *C. albicans* YHT network. Each method searches for different types of control strategies, with slightly different control objectives (Table 3). Nevertheless, across the analyses, some common key driver nodes and interactions emerged. Nrg1@HAGs, for example, was identified as a key node in all 4 control methods. In FVS control all but two feedback loops were broken by controlling Nrg1@HAGs (Figure S1). Nrg1@HAGs was a participant in almost all predicted stable motif control sequences (Figure 5). Further, simulations found that constitutive activity of Nrg1@HAGs could prevent the YHT, while constitutive inactivity of Nrg1@HAGs eliminated the yeast attractor. Lastly, more than half of the algebraic edge control predictions involved constitutive activity or silencing of edges involving Nrg1@HAGs or NRG1_T, including the interventions that led to the lowest total effect (Table 4). These control results agree with the well-known central gatekeeping role that Nrg1 plays in regulating the YHT [7].

Our edge control results predict effective parsimonious interventions that target interactions as opposed to nodes. For example, we predict that intervening in the acetylation properties of the promoter regions of HAGs in such a way to decrease the ability of Nrg1 to bind there would decrease the YHT. Such intervention offers a potentially more practical alternative than genetic deletion of *NRG1*. We predict that disabling the capacity of Brg1 to recruit HDACs, or disabling the capacity of HDACs to block HATs, would also disable the YHT.

While one may view these control predictions as possible targets for externally applied perturbations, they also reveal much about how the system may control its own repertoire of behaviors in different environmental conditions. Our analysis reveals the importance of positive feedback loops that rely on mutual inhibition; these feedback loops form conditionally stable motifs that can lock-in and restrict the system’s repertoire. The other regulatory motif important in this system is the incoherent feed-forward loop, which underlies both windows of opportunity discussed earlier. The most unexpected result concerns the incoherent feedback-mediated role of HATs in the YHT. First, HATs are required for hyphal_initiation [17], whereas they must be degraded for sustained hyphal maintenance [10]. The exact mechanisms mediating HAT activity, and their timing, requires further study.

There remain some experimental observations our model does not recapitulate, revealing gaps in knowledge regarding regulation of the YHT. For example, our current model does not recapitulate certain experimental observations pertaining to Brg1’s ability to recruit HDACs and to regulate HAGs, potentially questioning Brg1’s primacy among multiple interacting transcription factors that co-regulate HAGs [29]. Future genetic epistasis experiments, for example, that combine deletion of *BRG1* with deletion of genes encoding these interacting regulatory partners, may shed new light on the hierarchical roles of these transcription factors in regulating HAGs. The discrepancy regarding the phenotype of Efg1 constitutive activation suggests that Efg1 may be the mediator of the environment’s effect on *NRG1* transcription. We hope that our model will lead to follow-up experiments that eliminate these gaps of knowledge. Ultimately, given the importance of the yeast to hyphal transition in modulating *C. albicans* virulence, understanding how cellular decisions are made to undergo this morphological transition will ultimately allow us to better understand how *C. albicans* causes disease in humans and how this process can be altered to prevent disease.

## Methods

### Update methods

The state of the system at time *t* is given by an n-dimensional vector *x*^*t*^, where n is the number of nodes (15 in this case). For each node *i* in the network, there is a corresponding Boolean function *f* _*i*_(*x*^*t*^) that specifies the regulation of node *i*. To analyze the network, we used two different types of stochastic update for the Boolean functions: general asynchronous, and stochastic propensity.

In general asynchronous updating, at each time step one node (*i*) is selected at random and its value is updated as

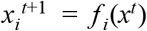

while all other nodes (*j* ≠ *i*) remain unchanged,

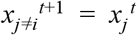

We also used the stochastic propensity framework described by Murrugarra et al [34]. Briefly, for each node *i*, we consider two parameters 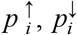 called activation and degradation propensities. 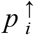 is the probability that in a situation when *f* _*i*_ calls for an update from 0 to 1 the state change happens (the state will remain 0 otherwise, with probability 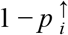) and 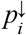 is the probability of an update from 1 to 0 according to *f* _*i*_. If applying *f* _*i*_ has no effect on σ_*i*_, then σ_*i*_ keeps its current value with probability 1. In summary,

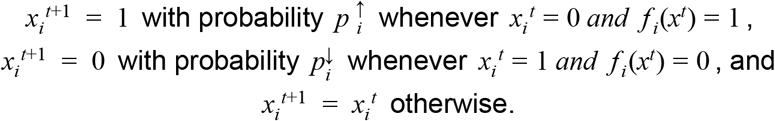

In the stochastic propensity framework, each node may have different propensities to change from 0 to 1 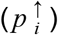 versus changing from 1 to 0 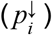, and these propensities may be different for different nodes. Unlike general asynchronous update, it is possible for multiple nodes to simultaneously change state in the stochastic propensity framework. Because of these differences, the timescales for simulations using the two frameworks are different. Due to the lack of information on the kinetic rates or timescales of the nodes in this network, we choose all the propensities to have value 0.5.

### State transition graph and attractors

Both Boolean update methods described above define a state transition graph (STG). Each possible state of the system corresponds to a node of the STG, and each directed edge indicates a possible next state after a single time step. In the general asynchronous update framework, only a single variable is updated each time step, so only states that differ by the value of a single variable can be connected in the STG. Conversely, in the stochastic propensity framework, multiple variables may simultaneously update within a single time step, and so distant nodes may be connected in the STG. This difference is partly responsible for the different timescales discussed above.

The state transition graph with all nodes and edges completely describes all possible dynamics of the system. Of special interest are the attractors of the system, which are individual states, or collections of multiple states, that have transition edges into them, but no transition edges out of them. In graph theoretical terms attractors correspond to terminal strongly connected components of the STG. Once the system enters an attractor, it cannot leave using the dynamics of the network alone. However, perturbations to the regulatory network, such as fixing node values or deleting edges, can change the underlying STG, possibly changing the set of attractors.

The distance between any two states *x, y* ∈ [0, 1]^*N*^ distance, defined as can be described using the Hamming

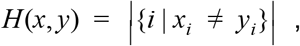

and which has the property that *H*(*x, y*) ∈ {0, 1, 2, …, *N*}.

Given any state *x*, the state *y* = (*f* _1_ (*x*), *f* _2_ (*x*), …, *f* _*N*_ (*x*)) describes the state achieved by applying each regulatory function *f* _*i*_(*x*) to each node *i* simultaneously. Let *d*_*x*_ = *H*(*x, y*) count the number of nodes whose values *can* change by applying one of the regulatory functions *f* _*i*_ (*x*). Each state *x* then has *d*_*x*_ possible transitions out under general asynchronous update. Specifically, the probability of transitioning between any pair of states *x* and *y* under general asynchronous update is

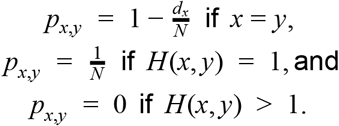

Conversely, under the stochastic propensity framework there are 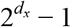 possible transitions out of each state *x* (the − 1 accounts for the case when none of the nodes that contribute to *d*_*x*_ update, which by definition is not a transition out of *x*). Each transition has probability 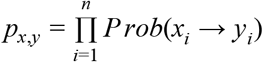 where

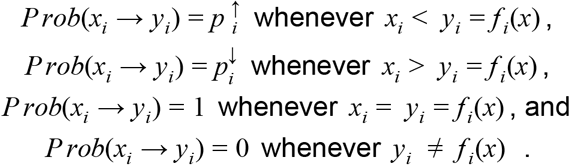

All transition edges present in the general asynchronous update STG are also present in the stochastic propensity STG, though they may have different probabilities. Conversely, the stochastic propensity STG may have many edges not present in the general asynchronous STG. Nevertheless, the two frameworks share many common dynamical behaviors.

If a state *x* is a point attractor (i.e., an attractor comprising a single state) of the system under general asynchronous update, then by definition of an attractor, *d*_*x*_ = 0. Thus there are 2^0^ − 1 = 0 transitions from *x* under stochastic propensity updates, indicating *x* is an attractor of the stochastic propensity system. Conversely, if *x* is a point attractor of the system under stochastic propensity updates, then 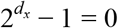, so *d*_*x*_ = 0. Thus, point attractors are preserved between both update orders. Indeed, point attractors are independent of the implementation of time [35].

In contrast, complex attractors are update-dependent [36], thus they are not guaranteed to be preserved between the two update methods. The *C. albicans* YHT network has five complex attractors under general asynchronous updates (Figure 2). Simulations of these attractors using stochastic propensity reveals that all nodes oscillating in the general asynchronous attractor also oscillate in the stochastic propensity attractor. Thus, for the YHT network, all attractors of each method are also attractors of the other.

### Stable motif analysis

Stable motifs correspond to positive feedback loops in the network that, once they achieve a certain state, become locked in. A stable motif succession diagram describes paths that may be taken once a given stable motif has locked in. Edges indicate subsequent stable motifs or conditionally stable motifs that may become locked in after the edge’s source stable motif is established. We created a stable motif succession network using the StableMotif python package [26]. Several branches of the stable-motif network contained redundant information that enabled simplifications. For example, the network has three inputs: pH, temperature, and Farnesol, each of which forms both ON and OFF stable motifs (e.g., pH=1 is a stable motif, as is pH=0). However, many of these lead to identical successions. Such redundancies were removed to derive a parsimonious version containing all the information (Figure 4).

### The total effect of deleting an edge

We implement the activation or deletion of a directed edge from node *s* to node *t* by replacing *x*_*s*_ in the regulatory function of node *t* by 1 or 0, respectively. To define the total effect of deleting an edge for stochastic propensity, we compute the number of changes in the state space before and after an edge deletion. If an update function *f* _*t*_ (*x*) is written in canalizing layers format [37], that is,

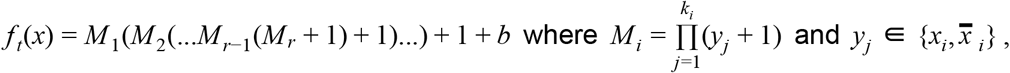

then the percentage of change from the initial state space upon the deletion of the edge from node *s* to node *t* is at most:

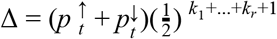

### Computation of the time to absorption

Consider the transition matrix of the Boolean network (as a Markov chain) in canonical form 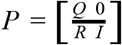, where 0 is the zero block matrix and I is the identity submatrix. The fundamental matrix N is defined as the inverse of (I-Q). That is, *N* = (*I* − *Q*) ^−1^. The time to absorption for a transient state j is defined as the expected number of steps before absorption and can be calculated as the sum of the jth row of N (see Theorem 11.5 of [38]).

### *C. albicans* Strains

The *C. albicans ume6* Δ/Δ strain (TF179) [39] and *hda1* Δ/Δ strain (gift from K. Kuchler) were constructed using the fusion PCR method described in [40]. The isogenic wildtype strain used for comparison was SN250 [41].

### Filamentation Assay

*C. albicans* cells were grown at 30°C on YPD agar plates for two days. Single colonies were picked and inoculated into YPD liquid medium and grown at 30°C overnight. Strains were inoculated from the overnight YPD culture into RPMI-1640 medium at pH=7.0 (with glutamine and phenol red and without bicarbonate, buffered with MOPS) at an OD600 = 0.2. RPMI-1640 cell cultures were incubated at 37°C for 90 minutes and imaged by light microscopy. A minimum of 70 cells were counted to quantify the percentage of cells categorized as yeast-form cells, transitional cells (including pseudohyphal cells), and true hyphal cells for each strain.

## Acknowledgments

We thank Karl Kuchler for the generous gift of the *hda1* Δ/Δ strain. We thank Jordan Rozum for his assistance with stable motif analysis.

## Supporting Information

**Figure S1.**
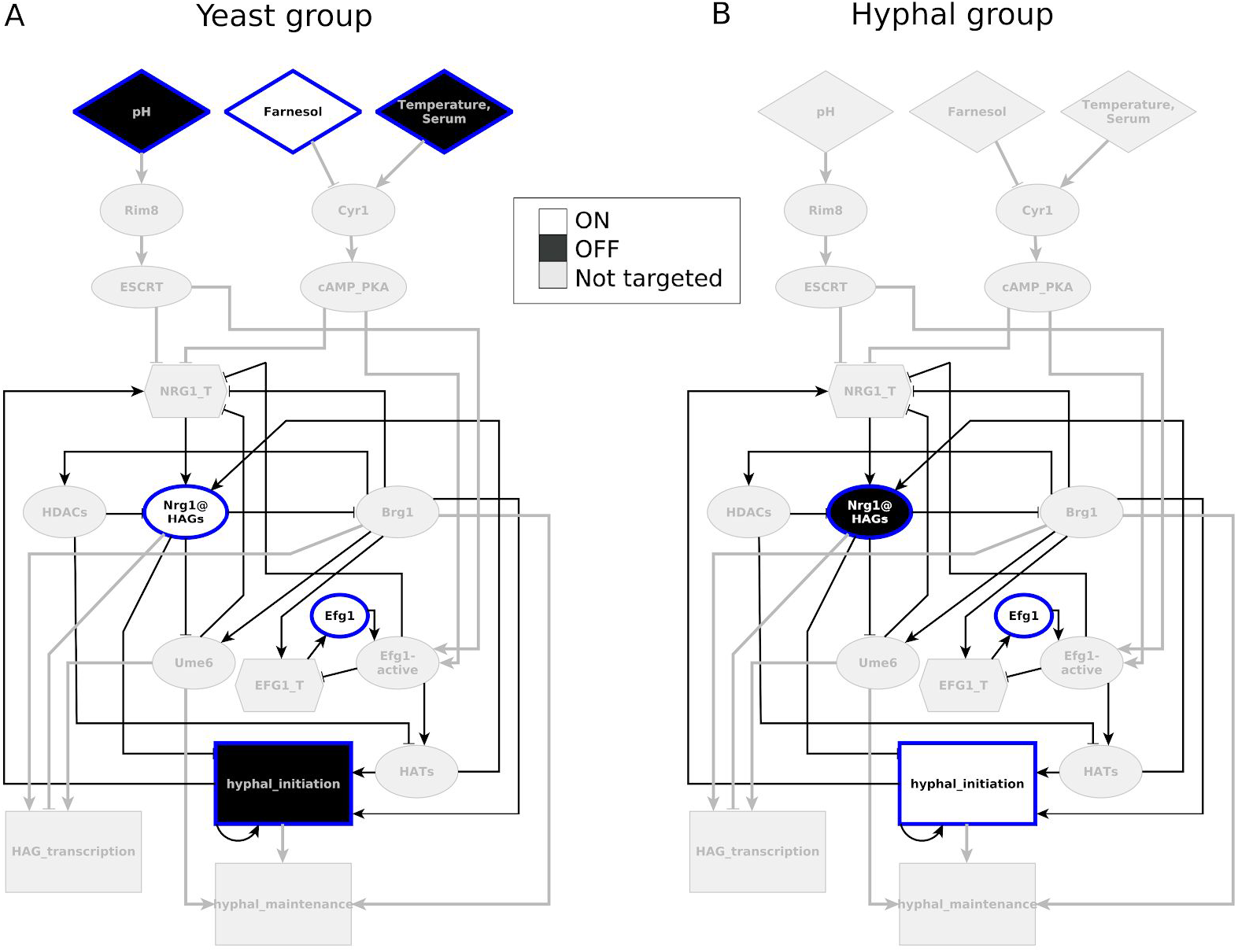
FVS control strategies to drive the system into a target attractor (group). Bold edges participate in feedback loops that are broken by controlling the values of the FVS. Nodes and edges that are irrelevant to feedback vertex control are shown in light grey, while nodes of the FVS are shown with a blue outline, and the nodes are colored based on the values they require for FVS control. (A) FVS control predicts fixing environmental conditions as pH = 0 and either Farnesol = 1 or Temperature = 0, and then fixing Nrg1@HAGs = Efg1 = 1 and hyphal_initiation = 0 will force the system into the yeast attractor. Instead of Efg1, EFG1_T or Efg1_active would also be suitable targets. (B) FVS control predicts fixing Nrg1@HAGs = 0, and Efg1 = hyphal_initiation = 1 will force the system into a hyphal attractor.

**Figure S2.**
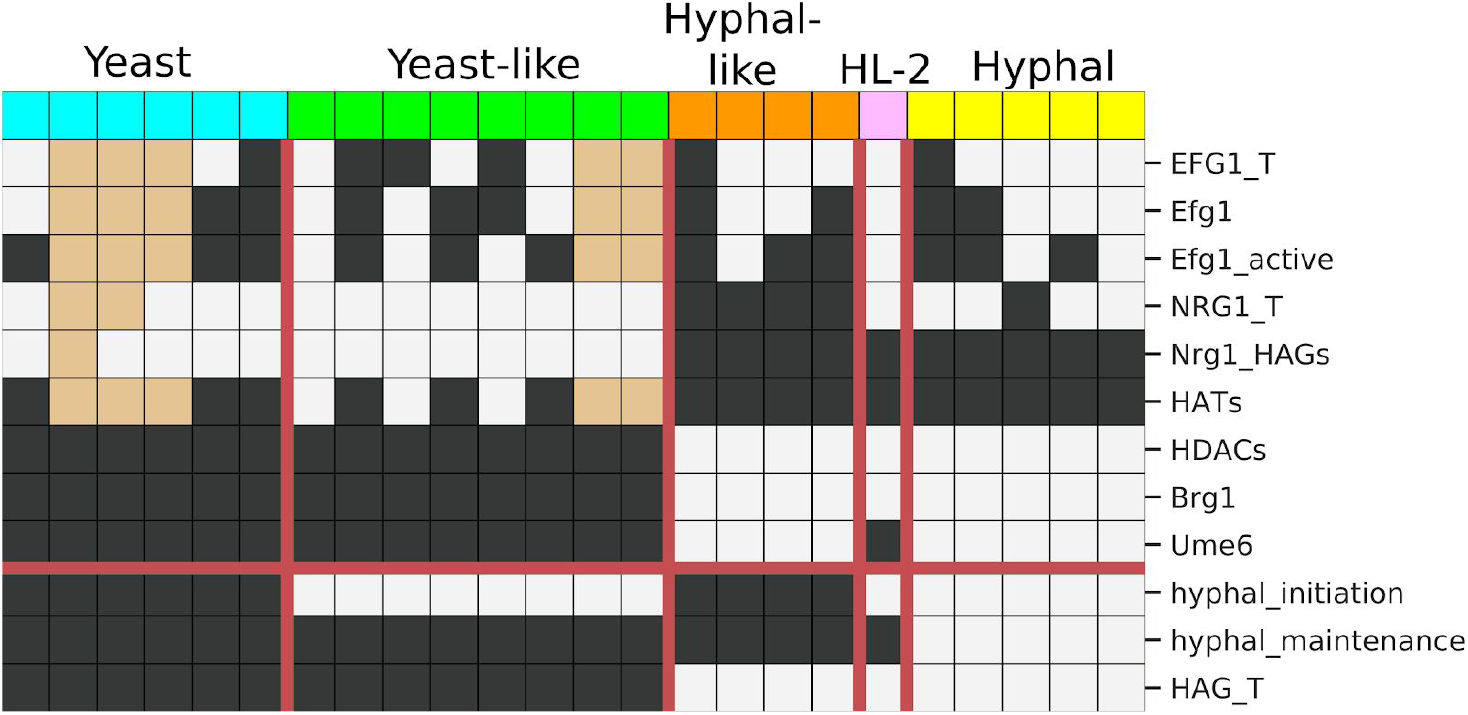
Single-node deletions or activations lead to new attractors. All attractors reached under YHT inducing conditions (pH = 1, Temperature = Farnesol = 0) are shown. Phenotype classification was performed as before, except the values of environment source nodes (pH, Farnesol, and Temperature), signaling intermediaries (Rim8, ESCRT, Cyr1, cAMP/PKA), and the individual perturbed node are ignored. One attractor emerged which did not fit the phenotype classifications defined previously, which we here call hyphal-like 2 (HL2). It corresponds to deletion of *UME6*, and has hyphal_initiation = 1, HAG_transcription = 1, and hyphal_maintenance = 0.

**Table S1.** Compilation of published experimental intervention results and comparison with the relevant model results. The first column describes the intervention. The second column indicates the environmental condition used in the experiments. The composition of the various media is the following: YPD (also denoted YEPD) medium: yeast extract-peptone dextrose, pH = 7 at the beginning of culture (it decreases as the yeast breaks down dextrose); B-medium: 0.67% yeast nitrogen base, 2% Na-succinate, pH = 6.5; RPMI-1640 supplemented with L-glutamine and buffered with morpholinepropanesulfonic acid (MOPS), pH=7.0; Spider medium: nutrient broth, mannitol, K2PO4, agar, pH = 7.2. Because all of these experiments start with a dilution of cells into fresh medium, the farnesol level is expected to be very low (equivalent with Farnesol = 0 in the model). The experimental conditions that lead to successful YHT in wildtype cells are shown in blue font; the rest are expected to be yeast-favoring environments. The third column summarizes the experimental result and the fifth column indicates the reference. The fourth column indicates the attractor repertoire of the model in the simulated intervention and environmental condition closest to the experiment. The model results that deviate from experimental observations are shown in red font. The rest of the model results are consistent with experimental observations.

| Expt | Environment | Phenotype(s) | Prediction | Ref(s) |
| --- | --- | --- | --- | --- |
| WT | YPD at 30°C | Yeast-form | 100% Y | [10] |
|  | YPD + 10% serum at 37°C | Extensive hyphal growth | 37% H, 51% HL, 12% YL | [10] |
| Efg1 = 0 | RPMI 1640 medium at 37°C | Deletion of *EFG1* leads to inability to filament, no biofilms, sparse monolayers of loosely attached elongated cells. | 100% Y | [42] |
|  | Spider medium at 37°C | Deletion of *EFG1* leads to inability to form biofilms and hyphae. |  | [29] |
| Efg1_active = 1 | B-Medium | Pseudohyphae | 100% Y | [30] |
|  | B-Medium +5% horse serum | Hyphae | 53% H, 27% HL, 20% YL | [30] |
| NRG1_T = 0 | YPD at 25°C | Pseudohyphae, HAGs are expressed. | 100% HL | [31] |
|  | YPD at 30°C | Hyphae |  | [32] |
|  | YPD + 20% serum at 37°C | Hyphae | 22% H, 78% HL | [31] |
| NRG1_T = 1 | Multiple hyphal-growth-inducing liquid media at 37°C | Inhibits hyphal growth in all conditions. | 100% Y | [43,44] |
| Brg1 = 0 | YPD + 10% serum at 37°C | Competent germ tube formation, defective hyphal elongation. | 100% Y | [27] |
| Brg1 = 1 | YEP Maltose medium at 25° C | Cells remained in yeast-form. | 100% HL | [27] |
|  | YEP Maltose medium at 37° C | Ectopic expression of *BRG1* sustains hyphal growth. | 97% HL, 3% H | [27] |
| HATs = 0 | YPD at 35°C | Deletion of *ESA1*, encoding a subunit of the HAT NuA4 complex blocked hyphal initiation. | 100% Y | [45] |
|  | YPD + 10% bovine serum at 37°C | Deletion of *YNG2*, encoding an active subunit of NuA4, diminished HAG transcription and formed few filaments. | 100% HL | [17] |
| HDACs = 0 | YPD + 10% serum at 37°C | Defective in sustained hyphal growth, Nrg1@HAGs increases after hyphal initiation. | 100% YL | [10] |
| Ume6 = 0 | YEPD + 10% fetal calf serum at 37 °C | Deletion of *UME6* leads to strong reduction in HAG transcription and significantly shorter filaments. | 42% HL2, 41% HL, 17% YL | [28] |
|  | YPD + 10% fetal bovine serum at 37 °C | Double deletion of *SSN6* and *RDP31* leads to decreased Ume6, and impaired filament extensions |  | [46] |
|  | YEPD at 30 °C | Yeast | 100% Y | [28] |
| Ume6=1 | YEPD at 30 °C | High level, constitutive expression of *UME6* leads to hyphal formation. Intermediate-level constitutive expression of *UME6* leads to a mixture of hyphae and pseudohyphae. | 100% HL | [18] |
| Delete edge from HDACs to HATs | YPD with 10% serum at 37°C | A constitutively acetylated Yng2 (active subunit of NuA4) could not sustain HAG transcription and hyphal elongation. | 100% YL | [10] |
| Rim8 = 1 or ESCRT =1 | Medium 199 at pH=4 and 29 °C | High Rim101 activity induced hyphal growth even at pH and temperature favoring the yeast-form. | 37% H, 51% HL, 12% YL | [47] |

### Text S1: Explanation of the regulatory functions of the model

**Rim8* = pH**

**ESCRT* = Rim8**

The pathway of sensing and responding to the environmental pH level is based on the Rim family of proteins. Several transmembrane proteins connected to intracellular Rim8 sense environmental pH. Neutral or alkaline pH (pH > 6) causes a signaling cascade through Rim8 that involves the endosomal sorting complex required for transport (ESCRT) and leads to cleavage of the transcription factor Rim101 [47]. To simplify we only include two elements, Rim8 and ESCRT, to describe activation of the full pathway in response to neutral/alkaline pH, which is abstracted to the Boolean state pH = 1.

**Cyr1* = Temperature and Serum and not Farnesol**

**cAMP_PKA* = Cyr1**

Cyclic AMP (cAMP) production by the adenylate cyclase Cyr1 and the subsequent activation of protein kinase A (PKA) is a major hyphal growth inducing pathway [15]. Serum promotes filamentation through the muramyl peptide, which binds to Cyr1 and stimulates its adenylate cyclase activity [13]. Incubation at 37 C leads to the activation of Cyr mediated by the molecular chaperone Hsp90 [14]. The quorum sensing molecule farnesol inhibits the activity of Cyr1 [12]. As the environments that consistently yield YHT include serum, have a temperature of 37 °C and a low cell density [4], we assume that Cyr1 is activated only if all three conditions are simultaneously satisfied. In the analysis of the model we merge the effects of serum and temperature into a single node, which we call “Temperature”.

**Efg1_T* = Brg1 or not Efg1_active**

**Efg1* = Efg1_T**

**Efg1_active* = (ESCRT or cAMP_PKA) and Efg1**

The Efg1 protein downregulates *EFG1* transcription in a negative self-regulation loop [9]. Brg1 can bind to the Efg1 promoter, it is unclear if it causes an expression change [29,48]. We include a positive effect from Brg1 to *EFG1* transcription. (This effect does not seem necessary for most findings.) We assume that either Brg1 or the absence of Efg1_active can maintain Efg1 transcription. We separate out the active form of the Efg1 protein (Efg1_active) from the generic Efg1 protein, which is translated from the *EFG1* transcript (Efg1_T). Efg1_active is induced in response to cAMP_PKA [49]. Based on the observation that the filamentation-inducing effect of constitutive activation of Rim101 is abolished by knocking out *EFG1* [47], we assume that the Rim pathway (ESCRT in the model) also leads to the activation of Efg1 (i.e. to Efg1_active).

This set of functions reproduces the observation that there is a reduction of Efg1 expression in response to YHT inducing stimuli [30]. These functions predict that Efg1 expression will be restored after Brg1 turns on.

**NRG1_T* = not Brg1 and not Ume6 and not (Efg1_active and (ESCRT or cAMP_PKA)) or hyphal_initiation**

Equivalently,

**NRG1_T* = not {Brg1 or Ume6 or [Efg1_active and (ESCRT or cAMP_PKA)]} or hyphal_initiation**

Signals that induce cAMP/PKA lead to the downregulation of *NRG1* [27]. We assume that pH acts similarly to inducers of cAMP/PKA in terms of how they affect the core network. Despite the downregulation of Efg1 expression following signals, Efg1 is required for downregulation of Nrg1 [10]. We implement this by considering that the active Efg1 is required, in collaboration with cAMP/PKA or pH signaling, for the downregulation of Nrg1.

Brg1 negatively influences the stability of the *NRG1* transcript by upregulating an antisense transcript [16]. Ume6 overexpression can repress Nrg1, and *UME6* KO reduces the downregulation of Nrg1 [28]. We include Ume6 as a sufficient inhibitor of NRG1_T.

We do not know the mechanism through which *NRG1* expression is restored following hyphal initiation, so we include a positive edge from hyphal_initiation to Nrg1_T. A possibility to consider in the future is that this effect is through Brg1, but time courses seem to suggest Brg1 increase and Nrg1 decrease are simultaneous [10].

**Nrg1@HAGs* = NRG1_T and (not HDACs or HATs)**

Equivalently,

**Nrg1@HAGs* = NRG1_T and not (HDACs and not HATs)**

The binding of Nrg1 to the promoter region of HAGs requires the expression of the NRG1 transcript and protein, and the correct chromatin state. HDACs (such as Hda1) deacetylate histones at the Nrg1 binding site of HAGs, preventing Nrg1 from binding at HAGs. However, the effect of HDACs is dependent on Hda1 first deacetylating the Yng2 subunit of NuA4 (a main contributor to the node HATs), leading to HATs degradation. After a YHT inducing signal, Nrg1@HAGs goes down because of Nrg1 expression downregulation. Even though Nrg1 expression can go back up, Nrg1@HAGs does not because of HDACs, which block Nrg1 from binding to the promoters of HAGs. If HATs are not degraded by HDACs, Nrg1 can return to its binding site [10] [27].

**Brg1* = not Nrg1@HAGs**

*BRG1* is one of the HAGs whose transcription Nrg1 blocks. Nrg1 and Brg1 form a mutual inhibitory loop [16]. Efg1, several other transcription factors encoded by HAGs, and Brg1 itself bind the promoter region of *BRG1 [29]*. As the strength of these effects is not known, we make Nrg1@HAGs the sole regulator of Brg1.

**HDACs* = Brg1**

Brg1 recruits the HDAC Hda1 to the promoters of HAGs [27].

**HATs* = Efg1_active and not HDACs**

HATs (such as NuA4) are induced by Efg1 [17]. The Yng2 subunit of NuA4 is deacetylated by Hda1, which causes its degradation [10]. We represent this effect as a negative regulation between HDACs and HATs.

**Ume6* = Brg1 and not Nrg1_HAGs**

Similarly to other HAGs, *UME6* transcription is regulated by Brg1 and HDACs [27]. We assume the effect of HDACs is through Nrg1@HAGs downregulation. Since HDAC KO disrupts *UME6* transcription [27], we assume an AND rule between Brg1 and “not Nrg1_HAGs”

**hyphal_initiation* = (HATs and Brg1 and not Nrg1_HAGs) or hyphal_initiation**

The recruitment of the NuA4 complex to the promoters of HAGs is required for nucleosomal H4 acetylation at the promoters during hyphal induction [17]. We implement this by assuming that HATs are required for hyphal initiation. We also assume that inactive Nrg1@HAGs and active Brg1 are necessary for hyphal initiation.

We assume that hyphal initiation is irreversible.

**HAG_transcription* = (Brg1 or Ume6) and not Nrg1@HAGs**

This phenotypic outcome node represents the process of transcription of hyphal-associated genes. Although its meaning is partially overlapping with the meaning of the process node “hyphal_initiation”, in our model it is possible to achieve HAG transcription while bypassing the standard mechanisms of hyphal initiation, by directly activating core TF drivers of the hyphal program. Having such a node allows the incorporation of the observation that Ume6 overexpression in yeast-favoring environments leads to transcription of HAGs and formation of pseudohyphae or hyphae [18]. We assume that the activity of either Brg1 or Ume6 and the inactivity of Nrg1@HAGs is needed for HAG transcription.

**hyphal_maintenance* = (Ume6 and not Nrg1_HAGs) and hyphal_initiation**

The phenotypic outcome node hyphal_maintenance expresses the hyphal development stage that follows hyphal initiation. We assume that hyphal initiation needs to have been completed for this phase to commence. The transcription factor Ume6 is expressed during hyphal elongation, and controls the level and duration of hyphal-specific genes and is important for hyphal elongation [10]. *UME6* levels are sufficient (even after HDA1 KO) for hyphal-maintenance [10,27]. Since Brg1 and HDAcs regulate *UME6* expression, we assume the effect of Brg1 and HDACs on hyphal maintenance is through *UME6*.

